# Signaling dynamics distinguish high and low priority neutrophil attractant receptors

**DOI:** 10.1101/2022.05.26.493652

**Authors:** Stefan M. Lundgren, Briana L. Rocha-Gregg, Emel Akdoǧan, Maya N. Mysore, Samuel L. Hayes, Sean R. Collins

**Affiliations:** Department of Microbiology and Molecular Genetics, University of California, Davis, Davis, CA, USA

## Abstract

Human neutrophils respond to multiple chemical attractants to guide their migration from the vasculature to sites of infection and injury, where they clear pathogens and amplify inflammation. To properly focus their responses during this complex navigation, neutrophils distinguish between attractants to modulate their responses. They prioritize pathogen and injury derived signals over long-range inflammatory signals secreted by host cells. Different attractants can also drive qualitatively different modes of migration. As receptors recognizing both classes of attractant couple to Gαi G-proteins, it remains unclear how downstream signaling pathways distinguish and prioritize between the inputs. Here, we use live-cell imaging to demonstrate that the responses differ in their signaling dynamics: low priority attractants cause transient responses, while responses to high priority attractants are sustained. We observe this difference in both primary neutrophils and differentiated HL-60 cells, for signaling outputs of calcium, a major regulator of secretion, and Cdc42, a primary regulator of polarity and cell steering. We find that the rapid attenuation of Cdc42 activation in response to LTB4 depends on the threonine 308 and serine 310 phosphorylation sites in the C-terminal tail of its receptor LTB4R, in a manner independent of endocytosis. Mutation of these residues to alanine impairs attractant prioritization, although it does not affect attractant-dependent differences in migration persistence. Our results indicate that distinct temporal regulation of shared signaling pathways distinguishes receptors and contributes to chemoattractant prioritization.

## INTRODUCTION

Neutrophils play central roles in inflammation and defense against microbial pathogens by responding rapidly to chemical signals and migrating to sites of injury or infection (*1, 2*). Once there, they mount cytotoxic responses including phagocytosis, degranulation, production of reactive oxygen species, and the release of anti-microbial extracellular traps (*3*). They also amplify inflammation by secreting factors that recruit additional immune cells (*4*). These activities are critical for effective immune responses, but their dysregulation is a major cause of tissue damage and chronic inflammation in autoimmune and inflammatory diseases (*5–7*).

Central to both productive and pathological inflammation, neutrophils navigate to target locations through chemotaxis, sensing gradients of chemical attractants with cell surface receptors. These receptors are almost uniformly G-protein coupled receptors (GPCRs) that couple to the Gαi family of G-proteins. While each of these receptors can drive chemotaxis, the behavioral responses to them are not identical. Different attractants can drive modes of migration that qualitatively differ in the directedness of their migratory paths (*8*), the gradient sensing strategies used (*9*), and the prioritization status when cells encounter competing gradients of different ligands (*10–12*).

Prioritization between attractants allows neutrophils to transition between guidance cues in complex environments where multiple chemoattractants often form overlapping gradients (*1*). Characteristically, host-secreted molecules such as the chemokine interleukin-8 (IL8) and the signaling lipid Leukotriene B4 (LTB4) act as “intermediary” signals, guiding neutrophils out of the vasculature and into the general vicinity of their target. However, these intermediary signals are ignored once neutrophils detect “end-target” signals near their ultimate destination. These injury or pathogen-derived signals, such as complement factor 5a (C5a) and formylated peptides (i.e., fMLF), are prioritized to focus neutrophils’ cytotoxic activities on the appropriate targets.

Differences in migration modes and prioritization status can be recapitulated in vitro, where cellular decisions depend almost entirely on the attractant identities and are largely independent of their relative concentrations (*8, 10–12*). However, the molecular mechanisms controlling these differences are not fully clear. In the case of prioritization, multiple mechanisms have been proposed, but it remains unclear whether prioritization is achieved through receptor-level regulation or crosstalk between downstream pathways (*10–17*).

In response to both intermediary and end-target attractants, receptor activation leads to a downstream signaling cascade driven by effectors of both Gαi and Gβγ subunits, including Rac, Cdc42, RhoA, calcium signaling, and reorganization of the actin cytoskeleton (Fig. 1A) (*18*). Notably, Cdc42 has been implicated as a primary regulator of cell polarity and cell-steering - making it an intriguing candidate for investigating potential differences in intermediary and end-target signaling inputs (*19–21*).

**Fig. 1.**
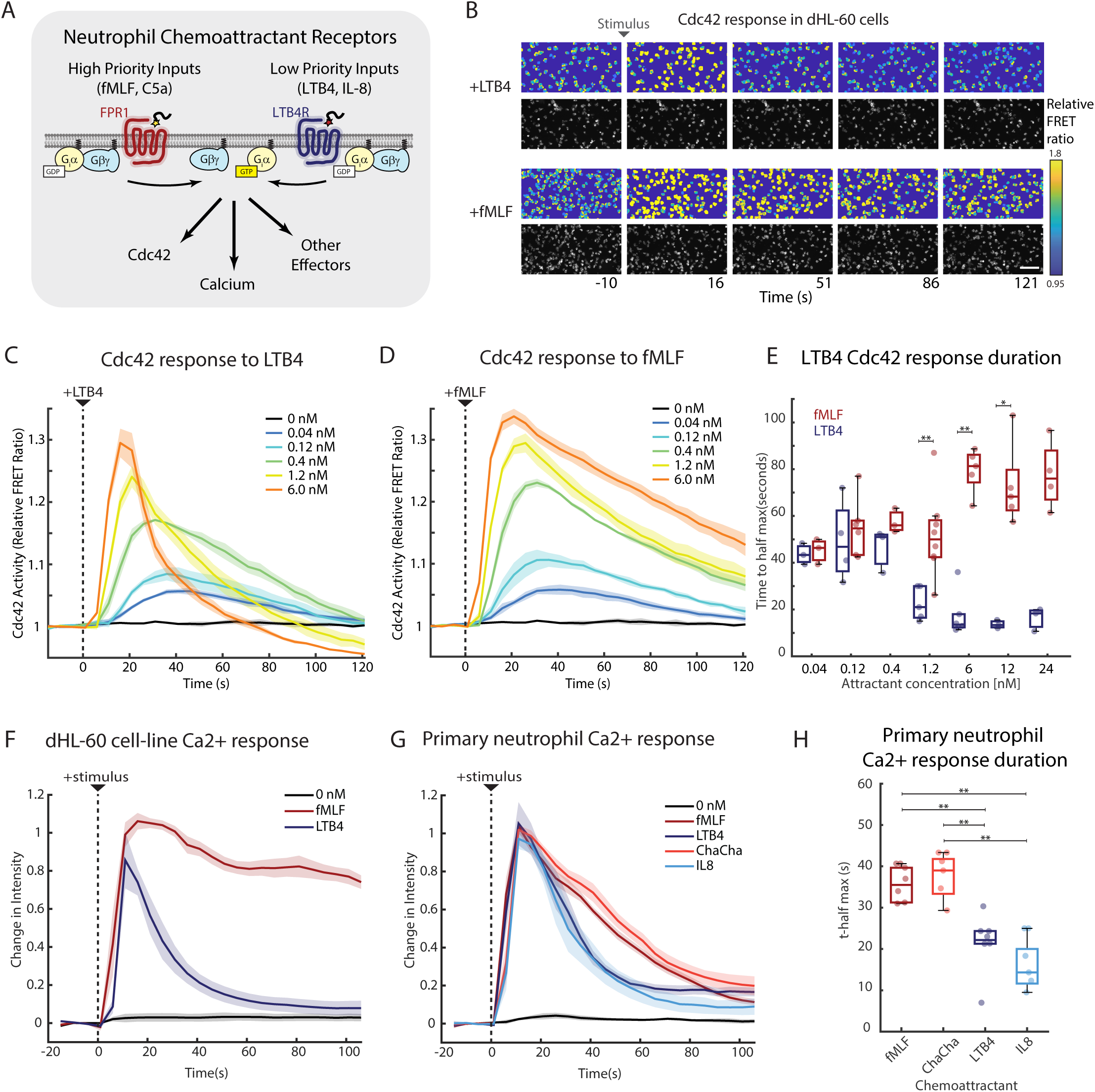
Low priority attractant receptors have faster signal attenuation than high priority receptors. (**A**) High and low priority receptor activation leads to a common downstream signaling cascade driven by effectors of Giα and Gβγ subunits. (**B**) Neutrophil-like differentiated HL-60 (dHL-60) cells expressing a Cdc42 FRET sensor were imaged at 5 second intervals before and after stimulation with 6 nM LTB4 or fMLF stimulus. Representative images show pseudo-colored relative FRET ratio above and grayscale of fluorescent intensity below. Scale bar, 100 μm. (**C, D**) Quantification of Cdc42 dynamics in response to stimulation with different concentrations of LTB4 or fMLF (n=3 for all conditions). (**E**) Boxplot showing the Cdc42 response durations calculated as the time from peak signal to half-maximal value. Dots indicate independent biological replicates. (**F, G**) Cytosolic calcium dynamics in response to stimulation with the indicated chemoattractants were measured by imaging dHL-60 or primary human neutrophils stained with Fluo-3 dye at 5 second intervals. Cells were treated with a saturating dose of “end-target” attractants fMLF (24 nM) or the C5aR agonist ChaCha peptide (1 μM), or “intermediary” attractants LTB4 or IL-8 (24 nM). Data was normalized according to the pre-stimulus baseline and the mean maximum signal for all conditions to account for staining variability. (**H**) Boxplot of the times to half max for primary neutrophil calcium responses. Dots indicate biological replicate measurements. For all panels, curves and shaded error regions represent the mean ± SEM over biological replicate measurements. Boxes in the boxplots indicate the 25th and 75th percentiles, with the center bar indicating the median, and whiskers indicating the range of the data aside from automatically determined outliers. P-values were calculated using the Mann-Whitney U-test, not shown for p-value > 0.05, * for p-value < 0.05, ** for p-value <0.01.

Here we found that the dynamics of signaling responses downstream of end-target and intermediary receptors differ, with end-target receptors generating a more sustained signaling response. We found that rapid attenuation of the response to LTB4 depends on the T308 and S310 phosphorylation sites in the C-terminal tail of the receptor LTB4R. Mutation of these two residues to alanines resulted in a sustained signaling response similar to that generated by the formyl peptide receptor, and it resulted in impaired prioritization of formyl peptide over LTB4. However, these mutations did not affect differences in migration persistence between these two attractants. Our experimental results are consistent with theoretical analysis demonstrating that homologous receptor desensitization could determine prioritization status (*15*). Our results suggest that signal output duration is a distinguishing feature between end-target and intermediary attractant receptors, and that these differences in signaling dynamics contribute to attractant prioritization.

## RESULTS

### Attractant receptors differ in output signaling dynamics

To understand how a cell can distinguish signals coming from different attractant receptors, we set out to determine whether the responses differ in the dynamics of activation of major downstream signaling pathways. We focused initially on fMLF and LTB4 as model end-target and intermediary attractants, and we used differentiated HL-60 (dHL-60) cells as a well-characterized model for human neutrophils. These cells can be differentiated into a neutrophil-like state, and we have previously characterized the broad similarity, as well as in gene expression between dHL-60 cells and primary human neutrophils (*22*).

We measured Cdc42 signaling activity in response to stimulation with a near-saturating dose (6 nM) of LTB4 or fMLF using a previously characterized, genetically encoded FRET-sensor (*19, 23*). For both chemoattractants, the population average Cdc42 response peaked within about 20 seconds, but the response to LTB4 attenuated much more rapidly (Fig. 1B, Movies S1, S2). After stimulation with LTB4, Cdc42 activity returned to baseline levels about 20-40 seconds after it reached its peak. Conversely, Cdc42 activity remained elevated for more than 2 minutes following fMLF stimulation (Fig. 1B, Movie S2).

We next asked whether the difference in signal attenuation arises from inherent properties of the corresponding receptors, FPR1 and LTB4R, or whether it could be explained by receptor sensitivity and the particular concentrations used. Thus, we performed a dose-response analysis.

At 0.04 nM, well below the reported Kd of each receptor (*24, 25*), the Cdc42 response was similar, with a low amplitude but relatively sustained duration. However, as LTB4 concentrations were increased, it induced stronger but increasingly transient Cdc42 responses (Fig. 1C). In contrast, as fMLF concentrations were increased, response amplitudes also increased, but the responses were sustained (Fig. 1D). For concentrations between 6 and 48 nM, the response patterns remained consistent, with rapid attenuation in response to LTB4 and sustained responses to fMLF (Fig. S1).

To quantify this more precisely, we measured the time required for the response to fall to its half-maximal level after its peak. As concentrations of LTB4 were increased from 0.4 to 6 nM, the time to half max decreased from ~40 seconds to ~10 seconds (Fig. 1E). For fMLF, the response duration became slightly longer as concentration was increased (Fig. 1E). Thus, saturating doses of LTB4 do not increase the duration of Cdc42 activation, nor do low doses of fMLF cause transient Cdc42 activity. These results indicate that LTB4R and FPR1 have qualitatively different signaling properties.

### Signal attenuation patterns are consistent across signaling outputs, attractants, and in primary human neutrophils

Given that LTB4R and FPR1 have distinct properties for regulating Cdc42, we wondered whether other shared downstream signaling pathways display corresponding differences in signal duration. In neutrophils, chemoattractant stimulation also induces the rapid release of calcium from the endoplasmic reticulum (*26*). The resulting cytosolic calcium increase regulates important neutrophil functions such as adhesion and degranulation. To determine whether LTB4 and fMLF stimulation have different effects on the duration of calcium signaling, we stained dHL-60 cells with the fluorescent calcium indicator dye Fluo-3 and imaged cells before and after the addition of 24 nM of LTB4 or fMLF. Indeed, we found that LTB4 induces a more transient elevation of cytosolic calcium than fMLF (Fig. 1F), indicating that the attractant-specific kinetics are transmitted to multiple signaling pathways. We note that the dynamics of calcium signals at the single-cell level are markedly different from Cdc42 dynamics, including pulsatile and oscillatory behavior (*27*). However, the consistent trend in signal attenuation at the population level suggests that the difference in signaling output of the receptors probably occurs upstream of both Cdc42 and calcium.

Next, we confirmed that differences in signaling properties between LTB4R and FPR1 are present in primary human neutrophils. While neutrophils are short-lived outside of circulation, the use of a calcium indicator dye allows measurement of signaling activity in freshly isolated neutrophils. We also extended our analysis of chemoattractants to include the ChaChapeptide (a C5α analog) that activates the end-target C5aR receptor (*28*) and the intermediary attractant interleukin-8 (IL-8) that activates CXCR1. Similar to our observations in dHL-60 cells, primary human neutrophils exhibit attractant-specific signaling dynamics. The intermediary attractants LTB4 and IL-8 induced a transient increase in cytosolic calcium concentration, while fMLF and ChaCha induced a longer lasting calcium signal (Fig. 1G). Again, we used the time to half-max as a measure of signal duration. We found no significant difference between attractants of the same class. However, for all comparisons between attractants of different classes, the end-target attractant had a significantly longer signal duration (p < 0.01) (Fig. 1H). We conclude that the attractant-specific differences in signaling dynamics in dHL-60 cells are representative of those in primary neutrophils, and that these differences may be a general feature of chemotaxis signaling that distinguishes classes of attractants.

### Chemoattractant-specific differences in signaling dynamics are evident in single cells

A limitation of population-level signaling studies is that population averages can mask differences in signaling activity only visible at the single-cell level (*29*). For example, the dose-dependent increases in amplitudes of Cdc42 activation (Figure 1C-D) could be due to increasing single-cell response amplitudes or due to an increase in the fraction of cells responding (with a consistent amplitude for all responding cells). Additionally, the persistence of Cdc42 activity in response to fMLF could be caused by slow attenuation at the single-cell level, or by asynchronous pulses of Cdc42 activity that, when averaged, give the appearance of a single, enduring signal. These alternate possible underlying behaviors would have important consequences for understanding signal integration in chemotaxis.

To distinguish between these possibilities, we performed single-cell analysis using data from the Cdc42 kinetics experiments described above. Overall, individual cell responses largely mirrored the population-level Cdc42 dynamics (Fig. 2A). In response to fMLF, single cells maintained persistent activation, rather than asynchronous pulses of activity, and at higher doses of LTB4, signals peaked and attenuated rapidly. However, we also observed apparent increases in the fraction of cells responding, which raised a question of whether responses were graded or “all-or-nothing” at the single cell level (Fig. 2A). Some signaling responses in chemotaxis show signatures of “excitable” behavior, including all-or-nothing responses (*30, 31*). In contrast, we have previously shown that Cdc42 responds in a graded manner to an optogenetic migration-directing GPCR (*32*).

**Fig. 2.**
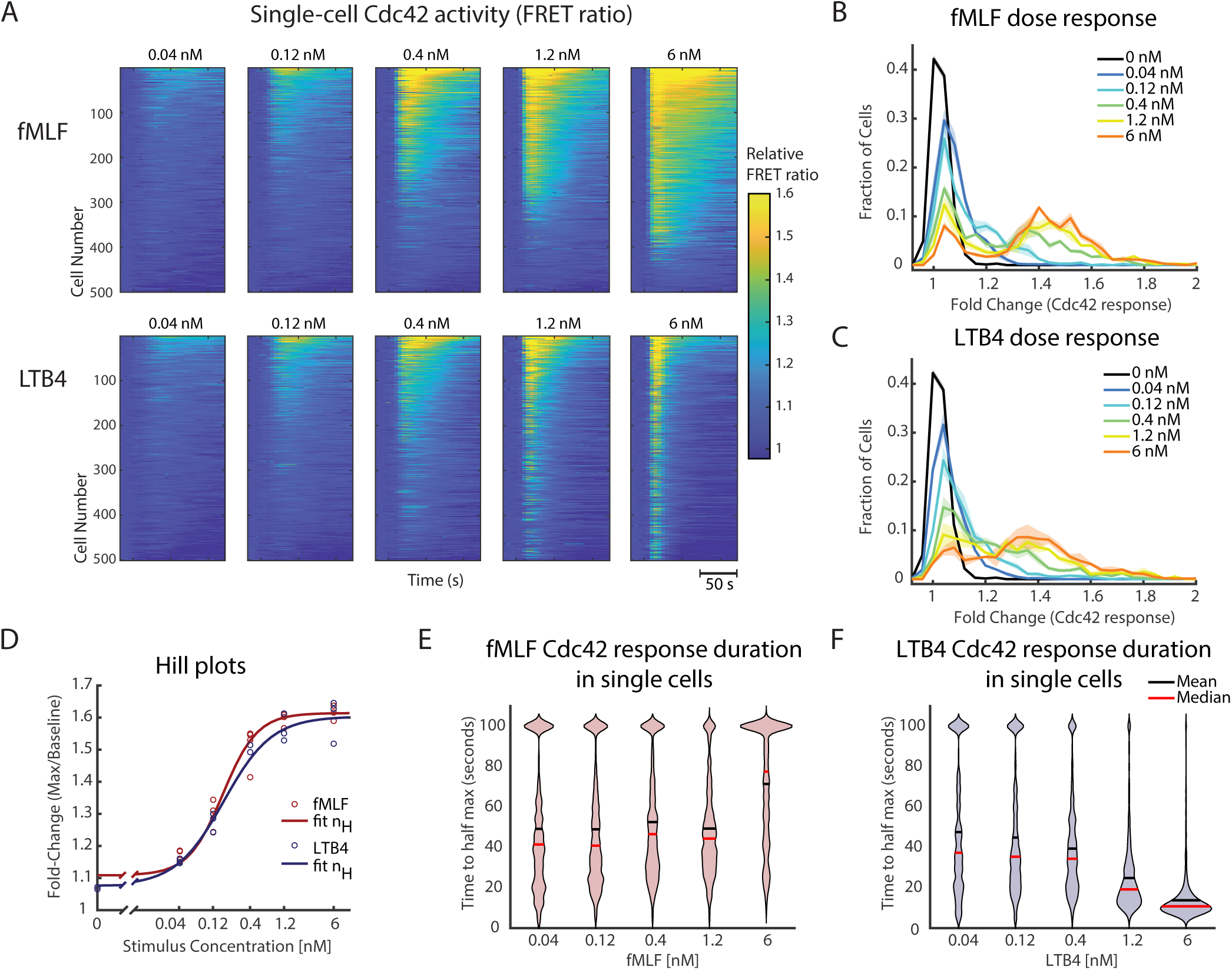
Single-cell kinetics are consistent with bulk analysis and reveal a graded response. Data presented in Fig. 2 are single cell information extracted from the experiments performed on dHL-60 cells in Fig. 1 (C, D). (**A**) Heat maps displaying Cdc42 FRET ratio over time on a single cell basis. Each concentration shows 500 cells selected as an even distribution of all cells analyzed. Data shown is arranged in descending order from top to bottom by the mean of the FRET ratio after stimulus. (**B, C**) Histograms displaying the frequency distributions of the single cell fold-change in Cdc42 FRET activity of baseline before stimulus compared to the maximum after stimulus. Curves and shaded error regions represent the mean ± SEM over biological replicate measurements. (**D**) Hill plots of Cdc42 response to chemoattractants. Dots indicate independent biological replicates, while the lines indicate the Hill-fit. Hill coefficient (nH) for fMLF = 1.80 ± 0.25 and for LTB4 = 1.28 ± 0.04. (**E, F**) Violin plots showing the distribution of Cdc42 response duration in response to fMLF (E) and LTB4 (F) among single cells.

To address this question more precisely, we computed histograms of the single-cell Cdc42 response amplitudes. For both attractants, the histograms showed some bimodal character, but there was a clear enrichment of intermediate responses at doses below 1.2 nM (Fig. 2B-C). The observation that the majority of responding cells in these conditions had sub-maximal amplitudes indicates that the response is indeed graded. The dose-dependent fraction of non-responding cells may arise in large part due to heterogeneity in receptor expression. We have previously measured expression of FPR1 in dHL-60 cells, and found that it can vary widely at the single cell level (*22*). Furthermore, we found that expression of an exogenous copy of LTB4R largely removed the population of nonresponding cells for stimuli of at least 0.6 nM LTB4 (Fig. S2). To analyze the ultrasensitivity of the response, accounting for the heterogeneity in receptor expression, we computed dose response curves considering the 90^th^ percentile response amplitude for each stimulus dose. We found that the response was moderately ultrasensitive, with hill coefficients between 1 and 2 (Fig. 2D). Analyses using different percentiles gave qualitatively similar results. These results are consistent with the presence of positive feedback in the Cdc42 signaling circuit (*32, 33*), and the observation of a graded response in which moderate attractant doses cause sub-maximal responses.

We further analyzed the signal attenuation by computing the time to half-max values for responses at the single-cell level. We only included responding cells, and we used an estimate of 100 seconds for cells whose signal had not yet dropped to the half-max value by the end of the timecourse. As in our population-level analysis, we found that the Cdc42 response attenuated rapidly for higher LTB4 doses, whereas the response to fMLF was sustained across the range of doses (Fig 2E-F). The dose-dependent nature of signal attenuation in response to LTB4 suggests that it is due to rapid negative regulation triggered downstream of LTB4R, acting on a timescale of roughly 10 seconds.

### Two phosphorylation sites control rapid signal attenuation for LTB4R

Given the attractant-specific, rapid attenuation of Cdc42 activity, we hypothesized that a negative regulator acts directly on the LTB4R receptor, shunting its signaling output. GPCRs contain numerous phosphorylation sites that play important roles in receptor-level regulation. In particular, phosphorylation of serine and threonine residues in the C-terminal tail is associated with ligand-induced receptor-desensitization (*34*). A recent study found that among multiple phosphorylation sites in LTB4R, two highly-conserved sites are phosphorylated in a ligand-dependent manner – T308 and S310 (Fig. 3A-B) (*35*). We focused on these residues as candidates for controlling signal attenuation and mutated the sites to alanine, both individually and in combination (Fig. 3B). We generated stable-cell lines exogenously expressing the mutant receptors in the HL-60 Cdc42 FRET-sensor background, reasoning that mutants affecting desensitization should be dominant, mediating their effects even in the presence of the endogenous wild-type copies of the gene. As a control, we generated a stable line that exogenously expresses the wild-type LTB4R under the same promoter.

**Fig. 3.**
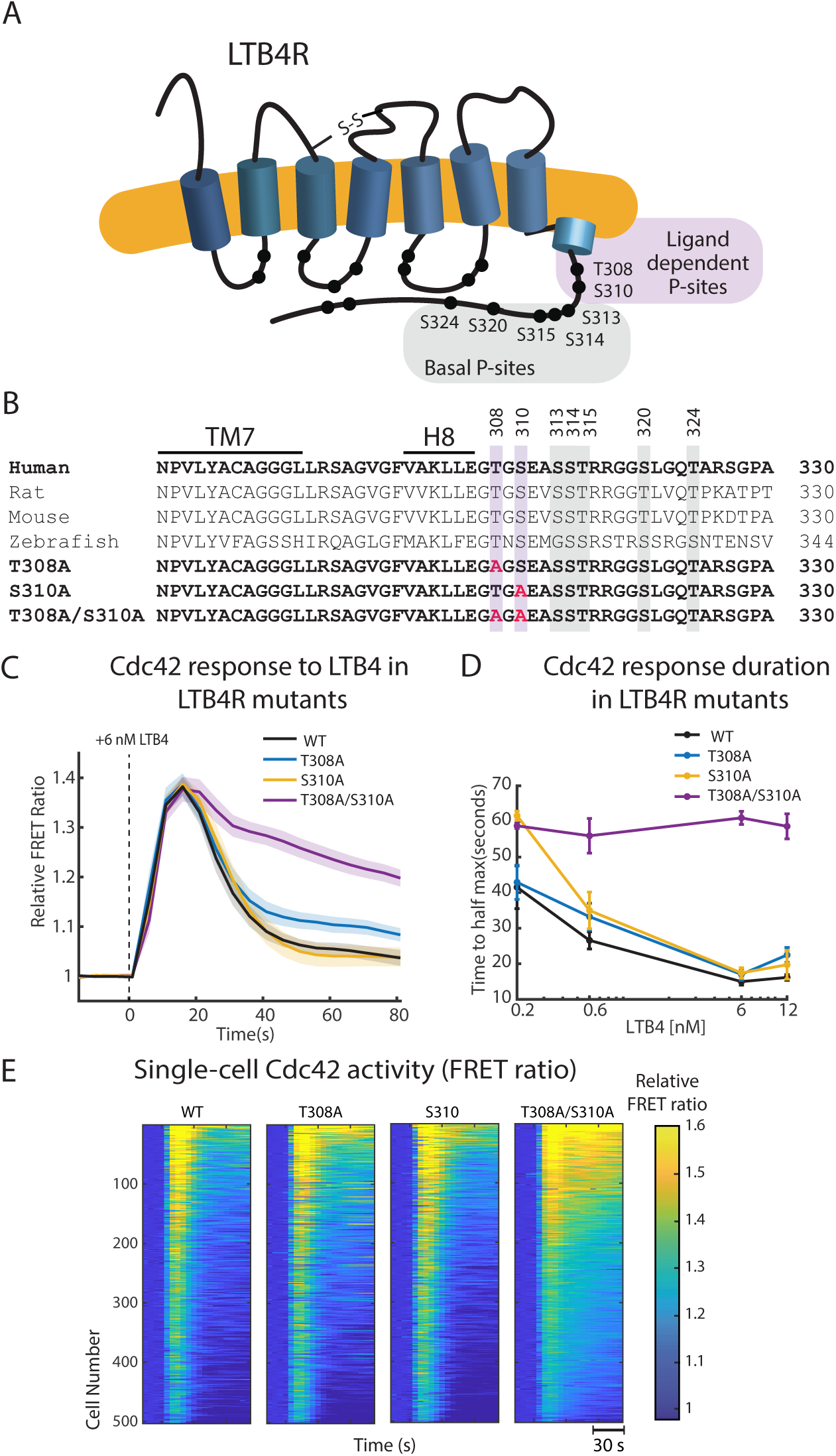
Rapid signal attenuation in response to LTB4 requires depends on T308 and S310 in the LTB4R1 receptor. (**A**) Schematic showing the LTB4R1 GPCR and phosphorylation sites on the C-terminal tail of the receptor. Phosphorylation sites are highlighted in purple are ligand dependent while those in grey are basally phosphorylated. (**B**) Amino acid sequence alignment of a section of LTB4R showing conservation across multiple metazoan species. Phosphorylation sites are highlighted using the color scheme in (A). Phosphorylation sites converted to alanines by site directed mutagenesis are shown in red. TM7= trans-membrane helix 7, H8 = helix 8. (**C**) Plots showing the relative Cdc42 FRET ratio of dHL-60 cells over-expressing versions of the LTB4 receptor under the same promoter. Images were acquired at 5 second intervals before and after 6 nM LTB4 stimulus. Curves and shaded error regions represent the mean ± SEM over at least five (n ≥ 5) biological replicate measurements. (**D**) Signal duration of the data in (C) was measured as time to half maximum as a dose response comparing mutant versions of the LTB4 receptor. (**E**) Single cell data was extracted from the experiment shown in (C, D). Single cell changes in Cdc42 activity are shown on heat maps, comparing different versions of the LTB4 receptor stimulated with 6 nM LTB4. 500 cells shown were selected as an even distribution from all cells.

While the individual T308A and S310A mutations had only small effects on the duration of Cdc42 activation, the double mutant receptors (T308A/S310A) eliminated the rapid signal attenuation, resulting in signaling dynamics similar to those induced by fMLF (Fig. 3C-E, Fig S3). The effects were consistent at the population level and in analysis of single cells. We quantified signal duration by measuring the time to half-max as before. Cells expressing wild-type, T308A, or S310A receptors showed dose-dependent decreases in signal duration, but cells expressing T308A/S310A receptors had approximately uniform signal duration for stimulus doses from 0.2 to 12 nM. To rule out potential confounding effects of receptor overexpression, we confirmed that LTB4R surface expression levels were comparable for all mutant and control lines using flow cytometry (Fig. S3). Therefore, the expression level alone cannot account for the signaling persistence observed in LTB4R-T308A/S310A cells. Our results indicate that both the T308 and S310 phosphorylation sites contribute to rapid LTB4R desensitization, and that either site alone is sufficient to induce desensitization.

### Signal attenuation is not caused by receptor endocytosis

GPCR phosphorylation plays a major role in regulating receptor endocytosis, and phosphorylation can induce desensitization through endocytosis or through endocytosis-independent mechanisms (*34*). Previous studies have found that LTB4R is resistant to endocytosis, which suggests that its desensitization is controlled separately (*36, 37*). We tested this idea by measuring the effects of the T308A and S310A mutations on ligand-induced receptor internalization by immunocytometry. Consistent with previous studies, we found that only 5-20% of wild-type LTB4R is internalized in response to 12 nM LTB4 stimulation (Fig. 4A-B). However, rather than blocking endocytosis, the T308A/S310A double mutation caused increased endocytosis. The effect was almost as strong for the T308A mutation alone, with S310A causing a small synergistic increase in endocytosis, but no measurable effect on its own (Fig. 4B, S4). Longer incubation with a higher dose of LTB4 (250 nM) induced higher rates of endocytosis, with about a 50% reduction in surface levels for wild-type LTB4R, but we found the same qualitative trends for the effects of the mutations (Fig. 4C, S4). Thus, T308 appears to be critical for limiting LTB4R internalization, with phosphorylation of this residue likely antagonizing internalization. Interestingly, T308 is separated by only two amino acids from the di-leucine motif in helix 8 that has previously been shown to be important for limiting LTB4R internalization (Fig. 3B) (*36*). These results demonstrate that ligand-induced desensitization of LTB4R occurs without large-scale endocytosis, and that in fact phosphorylation of T308 and S310 may have opposite effects on desensitization and endocytosis.

**Fig. 4.**
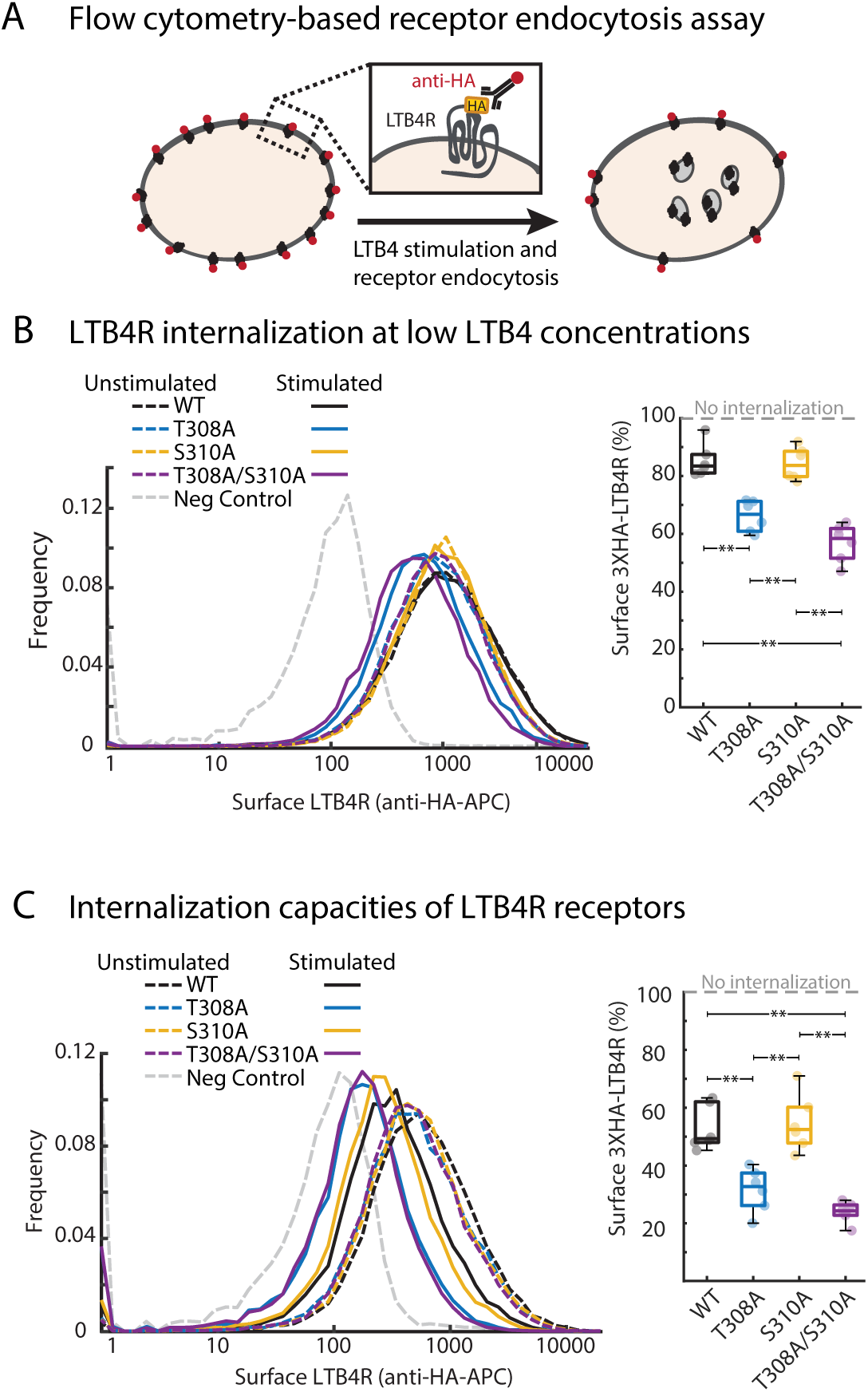
The T308 and S310 phosphorylation sites limit LTB4 receptor endocytosis. (**A**) A schematic depicting detection of LTB4R endocytosis measured by loss of fluorescent signal. Versions of the LTB4R receptor tagged with 3xHA (hemagglutinin-tag) were overexpressed in dHL-60 cells for detection on the cell surface by anti-HA antibodies conjugated to allophycocyanin (APC). Comparison of cells stained without LTB4 stimulation to cells with LTB4 stimulation determines percentage of receptor internalized. APC signal (LTB4R on the cell surface) was measured by flow cytometry. (**B**) Histograms showing LTB4R cell surface levels. Cells were either unstimulated or stimulated with 12 nM LTB4 for 10 minutes prior to staining with anti-HA antibodies (left). Boxplot (right) quantifying receptor internalization of LTB4R. 100% is defined as the surface LTB4R level in unstimulated cells and defines no internalization of the receptor. Representative of 5 independent biological replicates (n = 5). (**C**) Data showing experiments as performed in (B) but were treated with 250 nM LTB4 for 30 minutes to obtain maximum internalization of the receptors. Representative of 5 independent biological replicates (n = 5). P-values were determined using the Mann-Whitney U-test, not shown for p-value > 0.05, ** for p-value <0.01.

### Mutations affecting LTB4R signal attenuation impair attractant prioritization

Computational studies suggested that differing rates of homologous receptor desensitization, such as we characterized for LTB4R versus FPR1, could provide a mechanism for chemoattractant prioritization (*15*). The fast-desensitizing response to LTB4 could drive efficient chemotaxis in the absence of competing signals, but the more sustained output of FPR1 would cause it to dominate in the context of competing gradients. We set out to test this hypothesis by measuring the directionality of dHL-60 cells migrating in competing fMLF and LTB4 gradients.

We developed a simple under agarose chemoattractant prioritization assay in which gradients are established by diffusion from agarose “reservoirs” loaded with either no attractant or a chosen attractant at opposite ends of each well in a 24-well imaging plate (Fig. 5A). Cells migrate under an additional layer of agarose that covers the majority of the well, with attractant gradients formed by diffusion. We then track cell movement over time and measure speed and directionality. Using this assay, we found that dHL-60 cell migrate efficiently towards the gradient source for isolated gradients of either fMLF or LTB4 (Fig. 5B, S5). Consistent with prior observations, we found that these cells prioritize fMLF over LTB4 in competing gradients (Fig. 5B, S5) (*38*). Like cells expressing wild-type LTB4R, cells expressing the T308A/S310A mutant chemotaxed efficiently to both fMLF and LTB4 in isolated gradients. However, they showed a significant defect in prioritization, migrating with random directionality in the competing gradients (Fig. 5B).

**Fig. 5.**
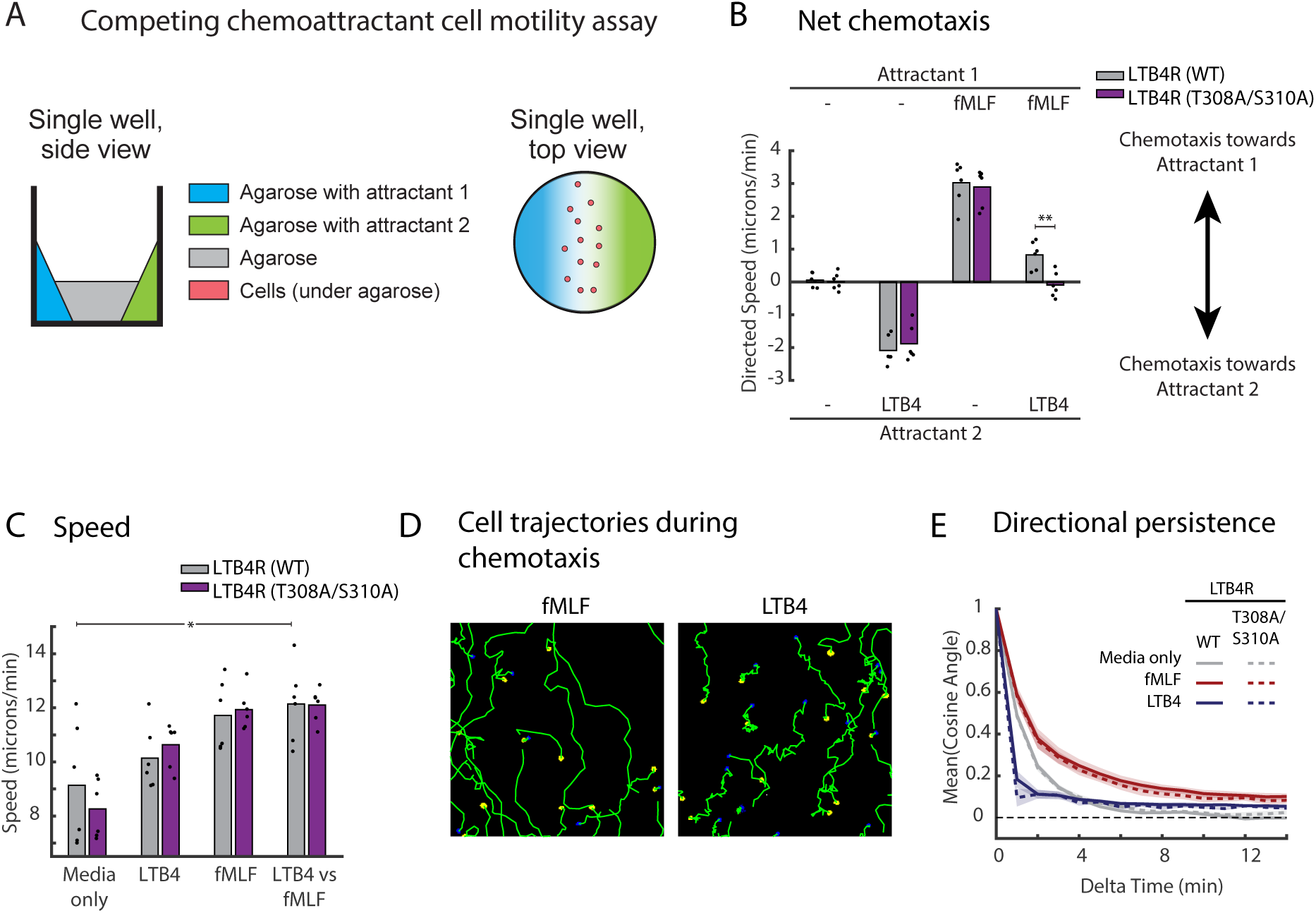
The T308 and S310 residues in LTB4R are necessary for attractant prioritization. (**A**) A diagram displaying the layout of a novel microscopy based competing attractant chemotaxis assay that was performed in a 24-well plate format. Cells move across the imaging plate surface under 1.5% agarose. Diffusion-based gradients were created by adding chemoattractant to “reservoirs” on one or both sides of the well and cell movement can be tracked over time by imaging (**B**) Chemotaxis of dHL-60 cells overexpressing LTB4R wild-type (wt) or LTB4R (T308A/S310A) cells was compared using the assay described in (A). Cells were stained with Hoechst stain and images were acquired for 60 minutes at a rate of 1 frame/minute. Net chemotaxis was defined as directed speed of cells toward fMLF (positive) or LTB4 (negative) and was measured as the rate of movement of the cell in the direction of the gradient source. Dots indicate independent biological replicates (n = 6). (**C**) Bar graph showing the average speed of cells in (B). (**D**) Images showing representative traces (green) of cell movement over 60 minutes. The initial (blue) image was merged with the final image (yellow) to show the starting and ending locations of the cells. (**E**) A plot showing the directional persistence of cell movement, defined as the mean cosine of the angle between a migrating cell’s movement direction at two different time points was measured as a function of the difference in time between the two measurements. Cells migrating in a straight line would have a mean cosine angle of 1. Data are presented as curves and shaded error regions that represent the mean ± SEM over biological replicate measurements. P-values were determined using the Mann-Whitney U-test, not shown for p-value > 0.05, * for p-value < 0.05, ** for p-value <0.01.

Analyzing these experiments, we also noted qualitative differences in the chemotaxis of dHL-60 cells to fMLF versus LTB4. First, the presence of fMLF seemed to trigger a larger increase in cell speed. We measured a significant increase in speed between the competing gradient condition and the no attractant condition (Fig. 5C). Although our other comparisons were not statistically significant, the higher speeds in the fMLF gradient suggest that fMLF may be largely responsible for the speed increase. Further experiments will be needed to determine how the attractant conditions differentially affect cell speed.

We also noticed that the movement trajectories of cells migrating in fMLF gradients were straighter, with fewer directional changes, than the trajectories of cells migrating in LTB4 gradients (Fig. 5D). We quantified the persistence of directionality in migrating cells by measuring the angle between a cell’s movement direction at timepoints separated by defined intervals in time. We found that directional persistence was increased in fMLF gradients but decreased in LTB4 gradients relative to random cell migration in the absence of attractant (Fig. 5E). We reasoned that this behavior might also result from differences in signaling dynamics, so we measured persistence in cells expressing the T308A/S310A mutant receptor. However, we found that the mutant receptor had no impact on the directional persistence (Fig. 5E, S5).

Our results indicate that the T308 and S310 phosphorylation sites in LTB4R contribute significantly to the ability of neutrophils to prioritize formyl peptides over LTB4 for chemotaxis. However, attractant-specific differences in other aspects of directed movement, such as directional persistence, must arise from different molecular or structural aspects of the receptors.

## DISCUSSION

Immune cells guide their migration using a family of GPCRs that activate Gαi, but the consequences of activating these receptors are not all the same. Distinctive features of these responses allow neutrophils to efficiently migrate to infection sites by prioritizing signals that emanate from sites of infection and limit their responses to prevent excessive damage (*4, 11, 39, 40*). However, the receptor elements and molecular mechanisms that determine these differences are not yet resolved, including how receptors detecting “end-target” ligands are prioritized over those detecting inflammatory signals secreted by host cells (*4*). Our results demonstrate that neutrophil responses to high and low priority chemoattractants differ qualitatively in the duration of signaling outputs. For LTB4R, this difference is determined by two phosphorylation sites, T308 and S310, in the C-terminal tail of the receptor, and these two sites are necessary for efficient prioritization of formyl peptide attractants over LTB4. Our results, combined with previous mathematical modeling (*15*), suggest a model in which the rapid desensitization of low priority attractant receptors causes their signals to be overpowered by the longer-lasting signaling output of high priority receptors (Fig. 6).

**Fig. 6.**
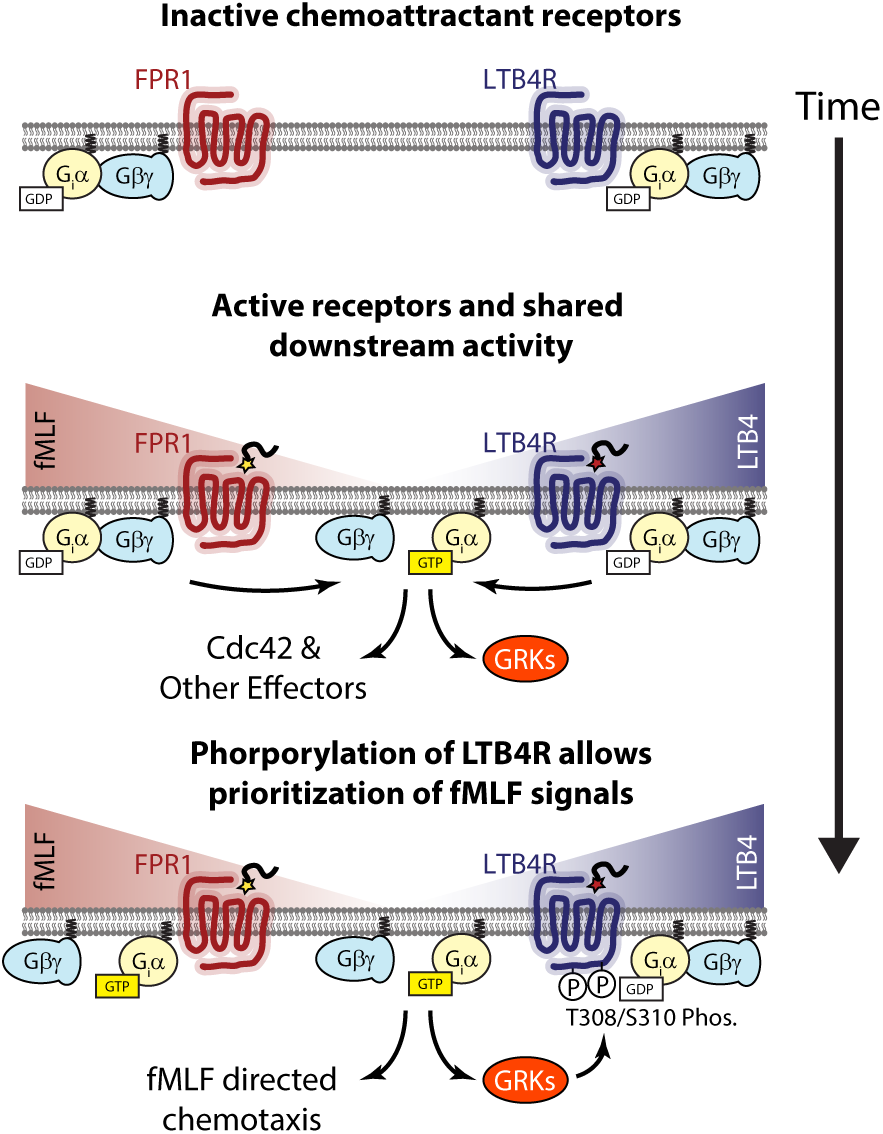
Shared signaling networks between chemoattractant receptors allow for prioritization of fMLF over LTB4 signals. A diagram displaying our model of signal transduction in response to multiple chemoattractant stimuli. In a resting state (top), inactive receptors are coupled with guanosine diphosphate (GDP) bound G-proteins. Exposure to chemoattractant gradients (middle) activates the receptors, leading to dissociation of G-proteins in their guanosine triphosphate (GTP) state and activity of other downstream signaling pathways. Following stimulus, G-protein coupled receptor kinases (GRKs) rapidly phosphorylate the LTB4R receptor (bottom) thereby switching the receptor off. The rapid desensitization for the LTB4R receptor allows the neutrophil to prioritize fMLF signals and the cell will move in an fMLF directed manor.

Receptor desensitization is broadly important for the function of immune cells, including neutrophils. It helps neutrophils self-limit swarming behavior at infection sites, it may be important for the reverse migration process that allows neutrophils to return to the vasculature from sites of inflammation, and we provide evidence here that it contributes to prioritization of attractants (*39, 40*). The GRK family of kinases plays a fundamental role in this desensitization, but the specific kinases and phosphorylation sites controlling receptor activity are not fully clear. In mice, GRK2 is required for the desensitization that limits LTB4-dependent neutrophil swarming (*39*), and a study using heterologous expression in COS-7 cells implicated T308 as a key residue for GRK6-mediated desensitization of LTB4R (*41*). Our results indicate that S310 plays an equivalently important role for LTB4R desensitization in human immune cells. It remains unclear which kinases regulate these sites, but a combination of GRK2, GRK6, and possibly additional kinases seems likely. In this more complex signaling network, it appears that T308 and S310 can each be phosphorylated rapidly, with either modification being sufficient to desensitize the receptor.

The differential homologous desensitization we characterized is likely to work in parallel with other mechanisms to achieve robust prioritization, and the full basis for prioritization may differ between pairs of receptors. There is evidence that downstream pathway crosstalk contributes to prioritization (*14*), although it is not clear how pathways are differentially engaged by the two receptor classes. There is also evidence that cross-desensitization contributes in some cases, where lower priority receptors are phosphorylated upon activation of higher priority receptors (*16, 42*). Even so, our calcium signaling results in primary neutrophils using IL-8 and the C5aR ligand ChaCha suggest that the receptor prioritization status may be linked more generally to the rate of homologous desensitization of the corresponding receptors.

Given the marked difference in signaling dynamics, it is interesting that neutrophils chemotax with similar efficiency to both LTB4 and fMLF (Fig. 5B, S3). A prior study provides a potential explanation, in that chemotaxis to low priority attractants relies largely on temporal sensing, rather than spatial measurement of gradients (*9*). Temporal sensing typically requires rapid signaling adaptation, allowing cells to sense changes in stimulus levels, rather than absolute levels (*43*). The rapid signal attenuation we see downstream of LTB4R could be well-suited for a role in temporal sensing by helping to reset downstream signaling on an appropriate timescale.

More generally, the behavioral responses triggered by attractant receptors differ in multiple ways. Aside from prioritization status, we found that LTB4 and fMLF drive qualitatively different migration patterns, with fMLF driving more persistent migration with straighter paths and fewer directional turns. Interestingly, the T308A/S310A mutations which blocked the characteristically rapid desensitization of LTB4R had no effect on the directional persistence of migration. This indicates that chemotactic GPCRs must have multiple sequence or structural differences that underlie attractant-specific features of responses.

## MATERIALS AND METHODS

### Cloning

Human *LTB4R* cDNA obtained from the Harvard plasmid repository (plasmid #HsCD00003892) was cloned to exogenously express *LTB4R* in HL-60 cell lines. Gibson assembly was used to clone *LTB4R* gene fragments into a lentiviral backbone containing the PGK-1 promoter and the hygromycin resistance gene downstream of an IRES sequence. In addition to the wild-type protein, T308A, S310A and T308A/S310A mutants were generated by site-directed mutagenesis using 20 cycle PCR reactions with Platinum-SuperFi DNA Polymerase (Invitrogen catalog #12359010). Construction of 3XHA versions of the human LTB4R sequence was performed using Gibson assembly. The 3XHA tag sequence was incorporated into primers. One Shot Stbl3 Chemically Competent *E. coli* (Invitrogen catalog #C737303) were transformed to amplify plasmid products. Plasmids were isolated from E. coli using plasmid preparation kits (Sigma) and sequence verified by Sanger sequencing (QuintaraBio).

### Cell culture

HL-60 cells were cultured in RPMI-1640 (Gibco, catalog #72-400-120) media with 9% heat-inactivated fetal bovine serum (FBS) (Sigma-Aldrich, catalog #F4135) and 100 U/ml penicillin-100 mg/ml streptomycin (Gibco, catalog # 15140163). The cell line used for these experiments is also known as PLB-985, which is a subline of HL-60. These cells were originally obtained as a gift from Dr. Orion Weiner’s lab. We have directly verified this genetic identity by analysis of SNPs (*22*). Cells were passaged every two to three days and maintained at a culture density between 1.0 × 10^5^ and 2.0 × 10^6^ cells/ml. Cells were differentiated into a neutrophil-like state by culturing an initial density of 2 × 10^5^ cells/ml in RPMI-1640 with 5% heat-inactivated FBS, 100 mg/ml, 1.3% DMSO, and 2% Nutridoma-CS (Roche, catalog #11363743001) (*22*). Cells were incubated this way for 6 days before use in experiments, at which point they were referred to as dHL-60 cells. HEK-293T cells (ATCC CRL-11268) were used for lentiviral production. Cells were cultured in high glucose Dulbecco’s modified Eagle’s medium (Sigma-Aldrich, D5671) that was supplemented with 9% FBS, 1% Glutamax (Gibco, catalog # 35050061), and 100 U/ml penicillin-100 mg/ml streptomycin. All cell lines were maintained in an incubator at 37°C and 5% CO_2_. For imaging experiments, a “modified L-15” imaging media (Leibovitz’s L-15 media lacking dye, riboflavin, and folic acid) (UC Davis Biological Media Services) was used to minimize media autofluorescence. Cell lines were regularly tested for mycoplasma. No mycoplasma contamination was detected for any cell line used in this work.

### Chemoattractants

Multiple chemoattractants were used throughout the experiment, including fMLF (Sigma-Aldrich catalog # 47729), LTB4 (Cayman Chemicals catalog # 20110), IL-8 (ThermoFisher Scientific, catalog # PHC0884), and ChaCha (Anaspec, catalog # AS-65121)

### Human primary neutrophil isolation

Ethical approval for the study of neutrophils from adult healthy controls was granted by the Institutional Review Board (IRB) from the University of California, Davis. All participants gave written, informed consent. Blood was collected by finger prick using a microlet lancing device. Neutrophils were isolated using negative selection with the EasySep Direct Human Neutrophil isolation kit (StemCell, catalog #19666), following the manufacturer’s instructions.

### Cell-line generation

Stable cell-lines were generated using 2nd generation lentiviral-mediated gene transfer. The packaging vector used was a gift from Dr. Lifeng Xu (pMD.G). The envelop vector was a gift from Dr. Peter Lewis (pCMV-dR8.2). HEK-293T cells were transfected in Opti-MEM (Gibco, catalog # 51985-034) using TransIT-2020 transfection reagent (VWR, catalog # 10767-014). Media was changed to DMEM containing 10% FBS after 12 hours. Supernatant was collected at 24 and 48 hours post media change and filtered (PES 0.45µm; catalog # 25-246). Lentivirus containing supernatant was concentrated 1:50 as described by the manufacturer (Origene, catalog # TR30026), flash frozen in liquid nitrogen and stored at −80°C. HL-60 cells were treated with 50 μl of the concentrated lentivirus and 2 mg/ml Polybrene (Sigma; catalog # H9268-5G). 18 hours later, infected cells were centrifuged (100G; 10 minutes) and resuspended in selection media (supplemented RPMI-1640 with 250 µg/ml Hygromycin). Exogenous, surface expression of receptors was confirmed via flow cytometry.

### Uniform chemoattractant stimulation

dHL-60 cells expressing a previously characterized CDC42-FRET sensor (*19, 23*) were resuspended in warmed modified L-15 media containing 2% FBS at a density of 500,000 cells/ml. To avoid washout, cells were plated on 96-well optical-glass bottomed imaging plates (Cellvis, catalog # P96-1.5H-N) coated with Poly-D-lysine (“PDL”, Sigma; catalog # P6407). In a sterile tissue-culture hood, 50 μl of 200 mg/ml PDL was added to each well and incubated at room temperature for 30 minutes. After incubation, wells were washed twice with 50 μl sterile DPBS (Life Technologies; catalog # 14190250) and dried thoroughly at 65°C for at least 30 minutes before 100 μl of cells were added (results in ~40-50k cells/well). Before imaging, plated cells were incubated at 37°C for 30 minutes to allow adherence. Time-lapse microscopy was performed using a Nikon Ti-E inverted microscope with a 20x (0.75 NA) Plan Apochromat objective. Images were taken at 5 second intervals. A flash of red light immediately preceding the 4th frame was used to signal the timing for the addition of 100 μl of chemoattractant or media by pipet. To achieve rapid mixing, chemoattractants were added to media at a 1:1 ratio. Images were captured using 2×2 binning. For FRET experiments, Dual Zyla-4.2-USB3 sCMOS cameras were used to capture CFP and YFP channels simultaneously.

### Calcium Assay

dHL-60 or primary neutrophils were centrifuged at 200g for 3 minutes and resuspended in sterile modified L-15 media containing 2% FBS, 2.5 mM probenecid (Life-Technologies: catalog # P36400) and 5 µM Fluo-3 (Life Technologies; catalog # F-14218). Cells were incubated at 37°C for 30 minutes before being centrifuged and resuspended in modified L-15 media containing 2% FBS and 2.5 mM probenecid. Uniform chemoattractant stimulation experiment was performed as described above. Quantification of fluorescent signal from Fluo-3 dye was performed using custom MATLAB scripts. Our image processing workflow included median background subtraction followed by calculation of the mean fluorescence in an image frame. Each independent biological replicate (n) consisted of the mean of three technical replicates. To account for day-to-day variability in Fluo-3 staining, data was first normalized to the mean of the first three frames and then to the mean of the maximum signal for each day.

### Flow-cytometry

Flow cytometry data acquired on a BD FACS Canto II flow cytometer (BD Biosciences, Franklin Lakes, NJ) was used to measure receptor internalization and expression of exogenous LTB4R. Flow cytometry data was analyzed using custom MATLAB scripts. Live cells were identified using gates based on FSC and SSC parameters. Surface-expression levels of LTB4R and mutant LTB4R in dHL-60 cells were assessed via immunostaining and flow-cytometry. Cells were harvested, washed with phosphate-buffered saline (PBS) and stained for 30 minutes on ice with a 1:50 dilution of AlexaFluor-647 conjugated anti-LTB4R antibody (R&D Systems; catalog #: FAB099R) in FACS buffer (0.5% bovine serum albumin and 0.05% sodium azide in PBS). After fluorescent labeling, the samples were washed with ice-cold FACS buffer and kept on ice until fluorescence was measured via flow cytometry.

### Receptor Internalization Assay

2×10^5^ dHL-60 cells in 100 μl were plated on a 96-well Costar assay plate (Corning). 100 μl of either complete medium (as a control) or 2X LTB4 in complete medium were added and mixed well. Cells were incubated at 37°C for either 10 min or 30 min. Samples were then transferred to ice, and centrifugations were performed at 4°C. We performed two washes after incubation using 150 μl FACS buffer (PBS with 5% heat-inactivated FBS and 0.01% sodium azide). The samples were then treated with 25 μl of APC-antiHA antibody with 1:75 dilution (~0.07 μg/well, BioLegend catalog # 901523) for 1 hour on ice in the dark. After two successive washes, the samples were resuspended in 200 μl of FACS buffer and analyzed by flow cytometry. Cells that were not stained were used to determine cellular auto-fluorescence, and cells that did not express an HA-tagged construct were used to access the background signal.

### Statistical analysis

All error bars and shaded error regions represent the standard error of the mean (±SEM) for the indicated number of independent biological replicates (n). Significance values were calculated using the nonparametric Mann-Whitney U-test (MATLAB’s ranksum function).

### Global and single-cell analysis of FRET images

FRET image pairs were analyzed using custom MATLAB scripts to register the images, subtract background, segment cells, and compute FRET ratios. Image registration was performed as described previously (*32*) to maximize alignment of cell edges in the two channels throughout the field of view, incorporating xy-translation, rotation, stretch in x and y dimensions, and second order terms to correct for optical aberrations. After registration, an approximation for the camera dark noise (100 intensity units) was subtracted, and then a shading correction was applied to correct for unequal illumination and light transmission across the field of view. Background subtraction was performed using the strategy previously described (*19*). Briefly, an initial objection detection and masking was performed using Otsu’s method to determine a threshold, and mask dilation to conservatively exclude cell pixels from the background computation. The background was then computed locally in 64×64 pixel regions as the median intensity of non-object pixels. The background was smoothed to avoid edge artifacts. This background calculation was performed separately for each channel, and the computed background images were subtracted. To generate cell masks, the two image channels were added together, sharpened using unsharp masking, log transformed, and thresholded. Masks were refined with image opening and the watershed algorithm to reduce noise and separate neighboring cells. All pixels not in the masks were set to NaN (not a number) to exclude them from further analysis. The donor and acceptor images were smoothed with gaussian filter of radius 1.5 pixels to reduce pixel noise. FRET ratio images were computed by diving the acceptor (YFP) image by the donor (CFP) image.

For bulk kinetic analyses, a single FRET ratio for the time point was computed as the sum of all acceptor intensities divided by the sum of all donor intensities, including only pixels in the mask. For single-cell analyses, cells were detected as distinct objects in the mask, excluding objects with an area below a minimum threshold of 150 pixels or above a maximum threshold of 900 pixels. The FRET ratio for each cell was computed as the sum of acceptor pixel intensities divided by the sum of donor pixel intensities for pixels in the object. Cells were tracked from frame to frame using a reciprocal nearest neighbor algorithm. Each independent biological replicate (n) consisted of the mean of three technical replicates.

Heatmaps of single cell signaling responses were generated using FRET ratios normalized to the mean of the baseline (3 frames before stimulus). A fraction of the cells are shown in the heatmaps. To make this selection, 167 cells were selected as an even distribution from each biological replicate. For visualization purposes, cells were sorted in descending order of FRET response to stimulus (defined as the mean FRET from frame 4 and later).

For generating histograms of signaling response, the fold change signal was calculated for each cell. We calculated fold change as the max FRET value after stimulus divided by the mean of the baseline. Using MATLAB’s ‘hist’ function, fold change values were sorted into 0.04 sized bins ranging from 0.92 to 2.0 and normalized to a frequency distribution. Each biological replicate (n) was counted as the mean of all values per day.

For the hill plots of the dose-response of Cdc42 signaling amplitudes, we computed the 90^th^ percentile fold-change among cells for each condition for each biological replicate. We used the 90^th^ percentile to estimate response characteristics for cells with high receptor expression, because we have found that heterogeneous receptor expression affects sensitivity to attractant. We have previously found that only approximately 70% of differentiated cells express high levels of FPR1 (*22*). Relatedly, we found that expressing an exogenous LTB4R largely eliminated the fraction of cells not responding to LTB4 at intermediate concentrations. Hill curves were fit using least squares regression and the Hill-Langmuir equation.

For violin plots of the Cdc42 response duration in responsive single cells. We defined responsive as meeting a 1.1 fold change above baseline in Cdc42 FRET activity. We computed the time to half-max (t-half) as the time from the max FRET ratio to the first value equal to or less than the value halfway between the max FRET ratio and the baseline FRET ratio. The max FRET ratio was chosen within the first 10 frames after stimulus to ensure the first response to stimulus was captured. If the FRET ratio did not decrease to half the max or less, time to half max was defined as 100 seconds (the max time possible from the median peak time to the end of image acquisition). The accumulation of cells at 100 seconds indicates cells that had a response duration longer than our imaging period. Violin plots were generated in MATLAB using kernel density estimation with bandwidth set to 2 (seconds).

### Under agarose chemotaxis and prioritization assay

To track cell movement in competing chemoattractant gradients, we developed an under agarose imaging assay with chemoattractant containing reservoirs of agarose on opposing sides of a well. 24-well optical-glass bottomed imaging plates (Cellvis, catalog # P24-1.5H-N) were coated with BSA and allowed to dry at 65°C. To prepare chemoattractant reservoirs, 3% low-melting point agarose (Bio Basic, catalog # AB0015) was prepared in modified L-15 media and cooled to 37°C, then mixed at a 1:1 ratio with a 2X concentration of chemoattractant in modified L-15 containing 4% FBS. 100 μl of the chemoattractant/agarose mixture was added to the imaging plate set on a stand holding the plate at a 70° angle above the counter surface. This allowed the agarose to solidify in the corner of the well. After 20 minutes, the plate was rotated, and chemoattractant/agarose was added to the opposing side of the well and allowed to solidify for 20 minutes at room temperature. The plate was set flat, and 1.5×10^4 cells stained with 1 μg/ml Hoechst (Life Technologies, catalog # H3570) in 5 μl modified L-15 with 10% FBS were dropped directly in the center of each well. After 5 minutes, 650 μl of 1.5% low melt agarose in modified L-15 containing 2% FBS was slowly added to each well, covering cells and reservoirs. The agarose was allowed set for 30 minutes at room temperature. The plate was sealed with an aluminum foil cover and incubated for 30 minutes at 37°C. Image acquisition over a 1-hour period at 1 frame/minute began immediately following the warming period.

### Tracking and statistics of cell movement

Cell movement was tracked and statistics were computed as described previously (*44*). Briefly, we used custom MATLAB scripts to identify cells, track them from frame to frame to assemble trajectories, and then compute statistics to measure multiple aspects of cell movement. Our image processing workflow included background subtraction, automated cell segmentation, and cell tracking. Cells were tracked from frame to frame by identifying the nearest neighbor in the latter frame for each cell in the prior frame (forward nearest neighbor) and the nearest neighbor in the prior frame for each cell in the latter frame (backward nearest neighbor), and requiring that the two methods agreed. Cell steps from frame to frame were linked to generate cell trajectories.

Next, we used the computed cell trajectories to calculate statistics. For every tracked cell step between adjacent frames, we computed a movement vector to determine distance moved and the angle of movement toward the chemoattractant gradient. An angle of 0° thus represented movement toward the center of the gradient, and an angle of 180° represented movement directly away from the center of the gradient. We applied a minimum distance moved threshold of 13μm (four pixels) for angle measurements to avoid noisy measurements for small movement steps.

We computed a directed movement length as the dot product between the movement vector and the optimal direction unit vector. From the aggregated cell step measurements, we computed mean cell speed as the mean of the movement distances divided by the corresponding time intervals. We computed mean directed movement as the mean of the directed movement distances divided by the corresponding time intervals. We computed angular bias as 90 minus the mean movement angle. Thus, an angular bias of zero corresponds to random direction relative to the gradient, an angular bias of 90 corresponds to maximal directionality toward the center of the gradient, and a negative angular bias corresponds to movement away from the center of the gradient. We computed the cosine of the angle between the direction of movement in the first 30 s, and the direction of movement in each subsequent frame-to-frame step. Only cells that moved at least 5 µm in the first 30 s step were included for analysis to capture only moving cells for which an initial direction could be determined accurately. We then computed the mean cosine value for each time point to determine the decay of directional persistence.

## Acknowledgments

We thank the Flow Cytometry Shared Resource at UC Davis and directors Bridget McLaughlin and Jonathan Van Dyke for their support and guidance. We also thank John Albeck, Marie Burns, Mark Huising, and the Collins’ lab members George Bell, Diana Sernas, Kwabena Badu-Nkansah, and Esther Rincón Gila for critical discussion throughout the project.

## Funding

National Institutes of Health grant DP2HD094656 (SRC)

Sidney Kimmel Foundation Kimmel Scholar Award (SRC)

National Institutes of Health IMSD grant R25GM056765 (BLRG)

Floyd and Mary Schwall Fellowship (BLRG)

National Institutes of Health Fellowship F31HL142150 (BLRG)

National Institutes of Health training grant T32GM007377 (SML, SLH)

National Institutes of Health P30CA093373

## Author contributions

Conceptualization: BLRG, SML, SRC

Methodology: SML, BLRG, SLH, MNM, EA, SRC

Investigation: SML, BLRG, SLH, MNM, EA, SRC

Formal analysis: SML, BLRG, EA, SRC

Software: BLRG, SML, SRC

Funding acquisition: SRC, BLRG

Writing – original draft: SML, BLRG, EA, SRC

## Data and materials availability

All of the processed data generated in this study are available in the main text or the supplementary materials. Due to the large size of the full data set, raw images are not included but are available upon reasonable request.

---

**Table S1.**
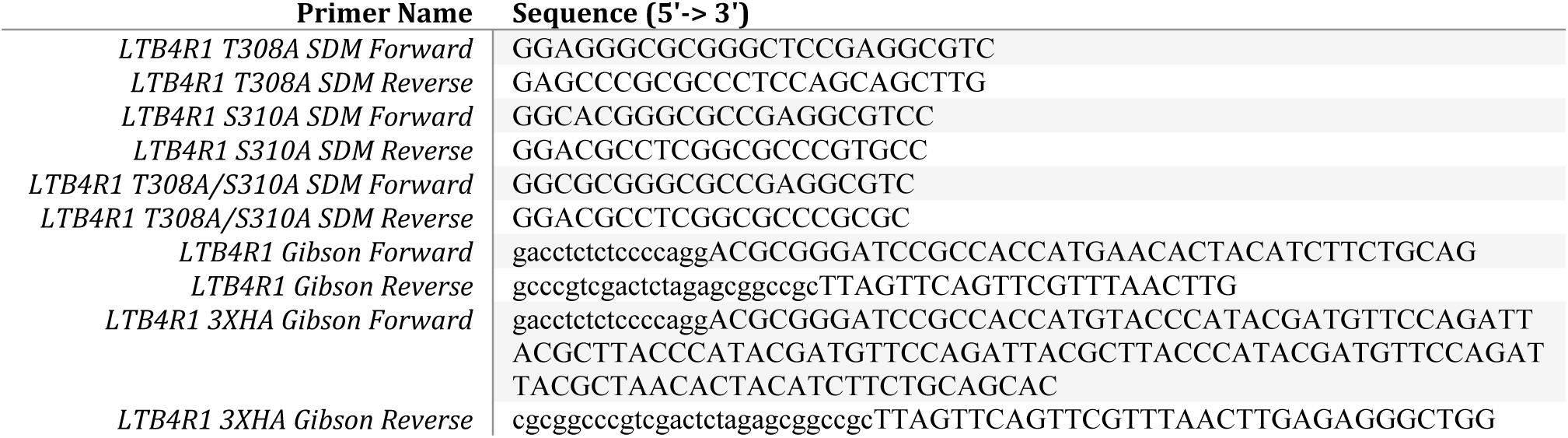
Primers used for generating plasmids.

## Supplementary Movie Legends

**Movie S1**. Cdc42 signaling response to stimulation with LTB4 in dHL-60 cells. Cdc42 signaling activity was measured with a FRET biosensor before and after stimulation with 6 nM LTB4 after frame 3. This movie shows the same experiment as in Figure 1B (top panels). Scale bar indicates 100 microns. The time of stimulus addition is defined as time zero.

**Movie S2**. Cdc42 signaling response to stimulation with fMLF in dHL-60 cells. Cdc42 signaling activity was measured with a FRET biosensor before and after stimulation with 6 nM fMLF after frame 3. This movie shows the same experiment as in Figure 1B (bottom panels). Scale bar indicates 100 microns. The time of stimulus addition is defined as time zero.

**Movie S3**. Movie of chemotaxis to fMLF with trajectories overlaid. dHL-60 cells were placed under agarose in a diffusion-generated fMLF gradient (fMLF source above the top of the movie frame). Movies show trajectories of cells as they moved over a 60 minute time period. Scale bar indicates 100 microns.

**Movie S4**. Movie of chemotaxis to LTB4 with trajectories overlaid. dHL-60 cells were placed under agarose in a diffusion-generated LTB4 gradient (LTB4 source below the bottom of the movie frame). Movies show trajectories of cells as they moved over a 60 minute time period. Scale bar indicates 100 microns.

