## Supplementary Figures for "Signaling dynamics distinguish high and low priority neutrophil attractant receptors"

Fig. S1

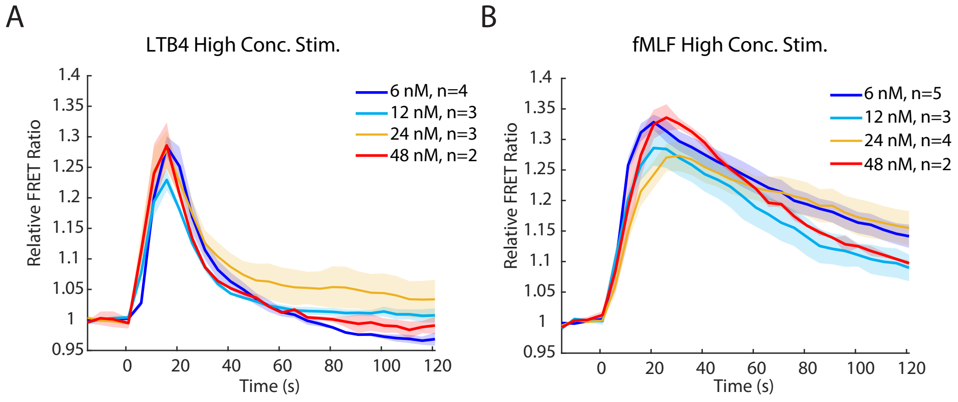


Fig. S1. Responses to LTB4 and fMLF are consistent at stimulus doses above 6 nM. (A, B) Quantification of Cdc42 dynamics in response to stimulation with high concentrations of LTB4 or fMLF. Curves and shaded error regions represent the mean ± SEM over biological replicate measurements.

Fig. S2

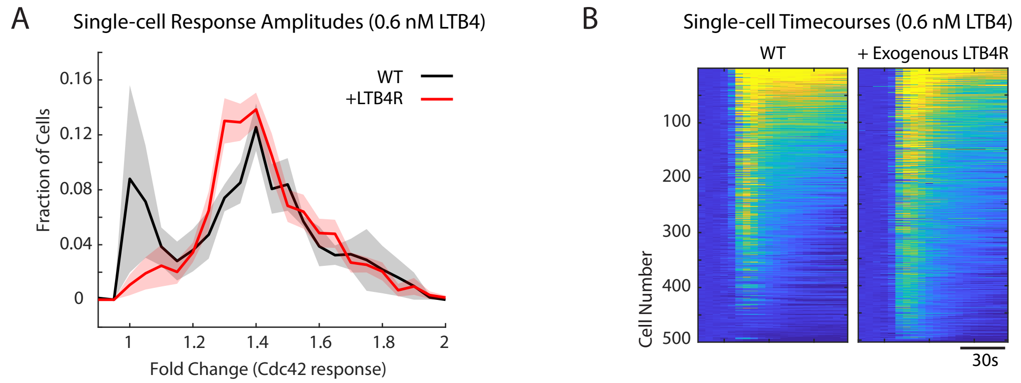


Fig. S2. Exogenous LTB4R expression yields homogenous response to LTB4. (A) Histogram displaying the frequency distributions of the single cell fold-change in Cdc42 FRET activity of baseline before stimulus compared to the maximum after stimulus. Curves and shaded error regions represent the mean ± SEM over n=4 biological replicate measurements. (B) Heat maps displaying change in Cdc42 FRET ratio in response to 0.6 nM LTB4 over time on a single cell basis. Data shown is arranged in descending order from top to bottom by the mean of the FRET ratio after stimulus.

Fig. S3

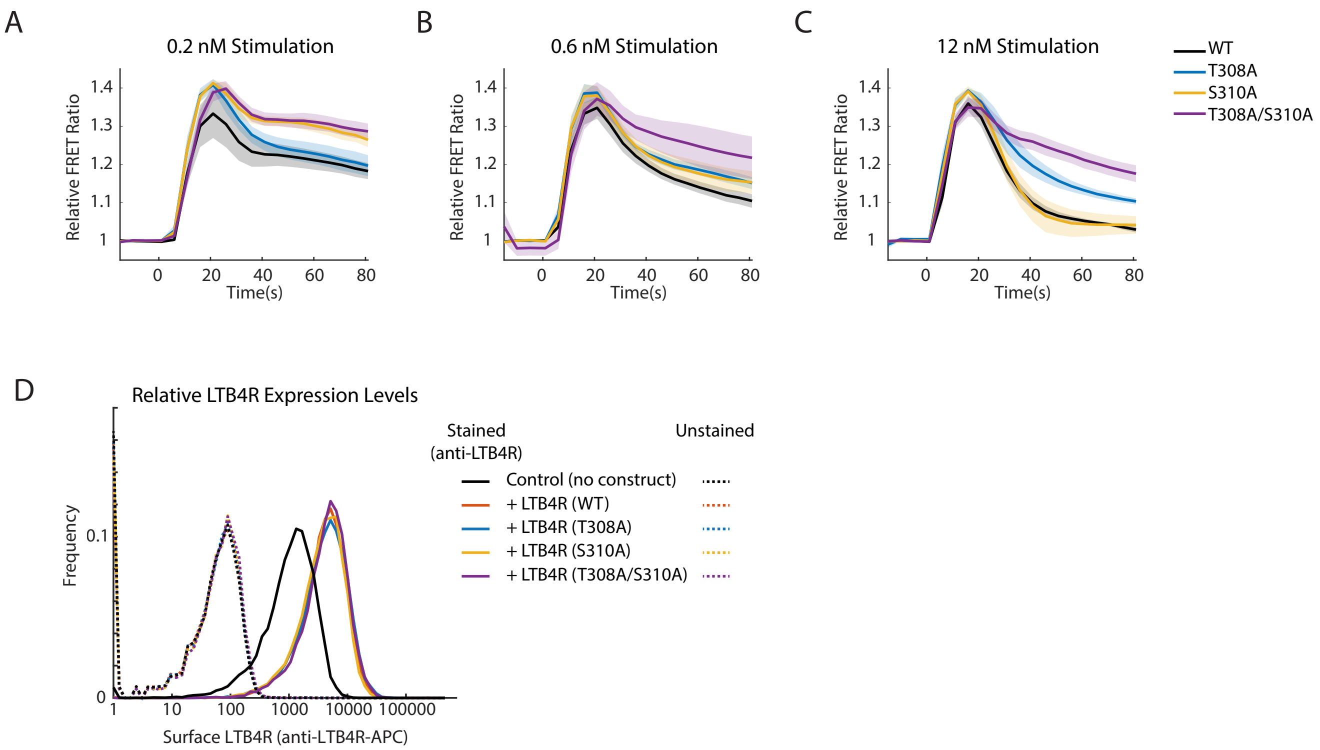


Fig. S3. Signaling and cell surface expression for LTB4R mutant receptors. (A-C) Plots with varying stimulus concentrations showing the relative Cdc42 FRET ratio of dHL-60 cells expressing the indicated LTB4R constructs. Images were acquired at 5 second intervals before and after the indicated concentration of LTB4 stimulus. Curves and shaded error regions represent the mean ± SEM over at least five ($n \geq 5$) biological replicate measurements. (D) Histograms showing quantification of LTB4R expression on the plasma membrane by flow cytometry. Conditions include cells with endogenous LTB4R only and cell lines expressing exogenous LTB4R constructs. Representative of three independent biological replicates.

Fig. S4

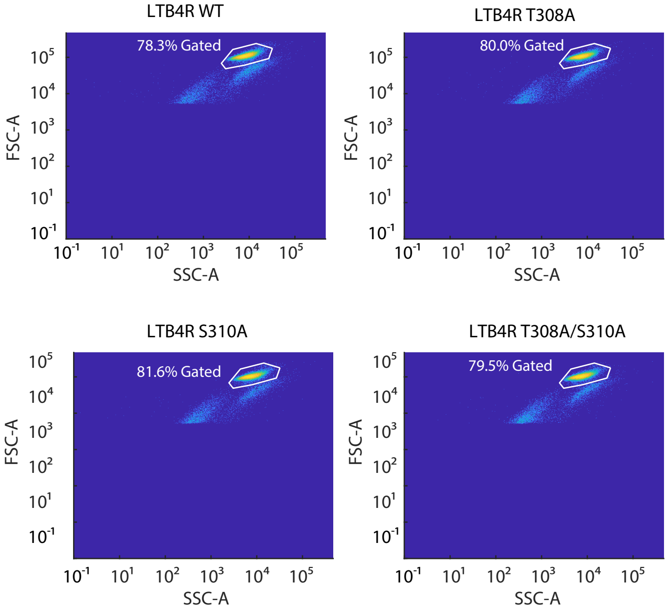


Fig. S4. Gates of live cells from a population of cells used for flow cytometry. Scatter plots displaying a representative gate for each of the indicated cell-lines.

Fig. S5

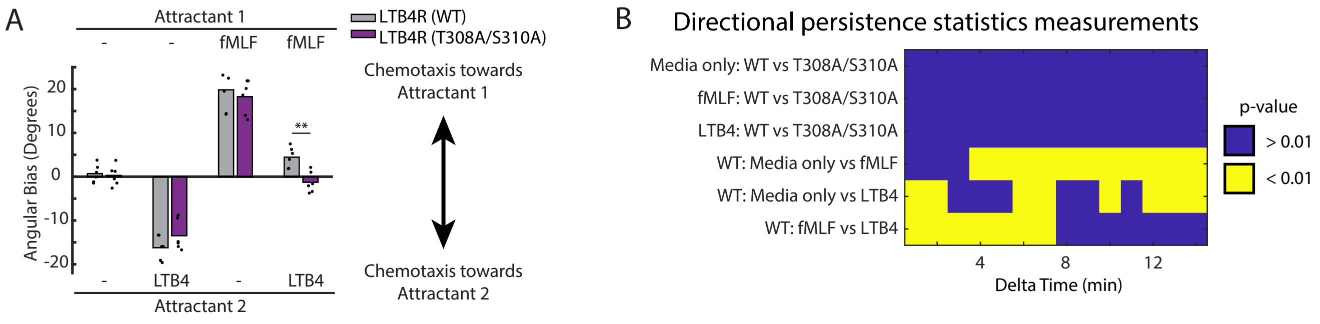


Fig. S5. Analysis of directionality in chemoattractant prioritization. (A) Quantification of the experiment shown in Fig 5B, measuring angular bias rather than directed speed. (B) Heatmap showing statistically significant comparisons of directional persistence between pairs of conditions. Significance was determined based on a false discovery rate threshold of 5%, which resulted in a maximum p-value threshold of 0.01 and a computed false discovery rate of 0.032. For each time interval (delta t), a p-value greater than the threshold is indicated with blue while a p-value less than the threshold is indicated in yellow.
